# Triple-helix potential of the mouse genome

**DOI:** 10.1101/2021.12.31.474609

**Authors:** Kaku Maekawa, Shintaro Yamada, Rahul Sharma, Jayanta Chaudhuri, Scott Keeney

## Abstract

Certain DNA sequences, including mirror-symmetric polypyrimidine/polypurine runs, are capable of folding into a triple-helix-containing non-B-form DNA structure called H-DNA. Such H-DNA-forming sequences are frequent in many eukaryotic genomes, including in mammals, and multiple lines of evidence indicate that these motifs are mutagenic and can impinge on DNA replication, transcription, and other aspects of genome function. In this study, we show that the triplex-forming potential of H-DNA motifs in the mouse genome can be evaluated using S1-sequencing (S1-seq), which uses the single-stranded DNA (ssDNA)-specific nuclease S1 to generate deep-sequencing libraries that report on the position of ssDNA throughout the genome. When S1-seq was applied to genomic DNA isolated from mouse testis cells and splenic B cells, we observed prominent clusters of S1-seq reads that appeared to be independent of endogenous double-strand breaks, that coincided with H-DNA motifs, and that correlated strongly with the triplex-forming potential of the motifs. Fine-scale patterns of S1-seq reads, including a pronounced strand asymmetry in favor of centrally-positioned reads on the pyrimidine-containing strand, suggested that this S1-seq signal is specific for one of the four possible isomers of H-DNA (H-y5). By leveraging the abundance and complexity of naturally occurring H-DNA motifs across the mouse genome, we further defined how polypyrimidine repeat length and the presence of repeat-interrupting substitutions modify the structure of H-DNA. This study provides a new approach for studying DNA secondary structure genome wide at high spatial resolution.

## Introduction

DNA sequence motifs with potential to form non-B-form DNA structures are enriched in noncoding regions of eukaryotic genomes (Bucher and Yagil, 1991; Behe, 1995). Several lines of evidence suggest that these motifs have biological impacts (Bacolla and Wells, 2009; Zhao et al., 2010; Kasinathan and Henikoff, 2018; Guiblet et al., 2021). However, their non-B DNA forming potentials are still not fully understood.

One such non-B-form DNA structure is H-DNA, which consists of an intramolecular triplex plus single-stranded DNA (ssDNA) regions (**Figure 1A**) (Mirkin and Frank-Kamenetskii, 1994). In H-DNA, the duplex from one half of the H-DNA motif forms Hoogsteen triplets with a “third strand” contributed by melting the duplex in the other half of the motif. Polypyrimidine/polypurine mirror repeat sequences are prominent examples of such H-DNA forming motifs. H-DNA contains three ssDNA regions that we will refer to as the central loop, orphan strand, and junction (**Figure 1A**). The central loop is a ssDNA hairpin loop between the triplex forming segments on the same strand that contributes the third strand to the triplex. The junction is a short ssDNA region between the triplex and the flanking duplex. And the orphan is the strand that is complementary to the central loop, the third strand, and the junction.

**Figure 1.**
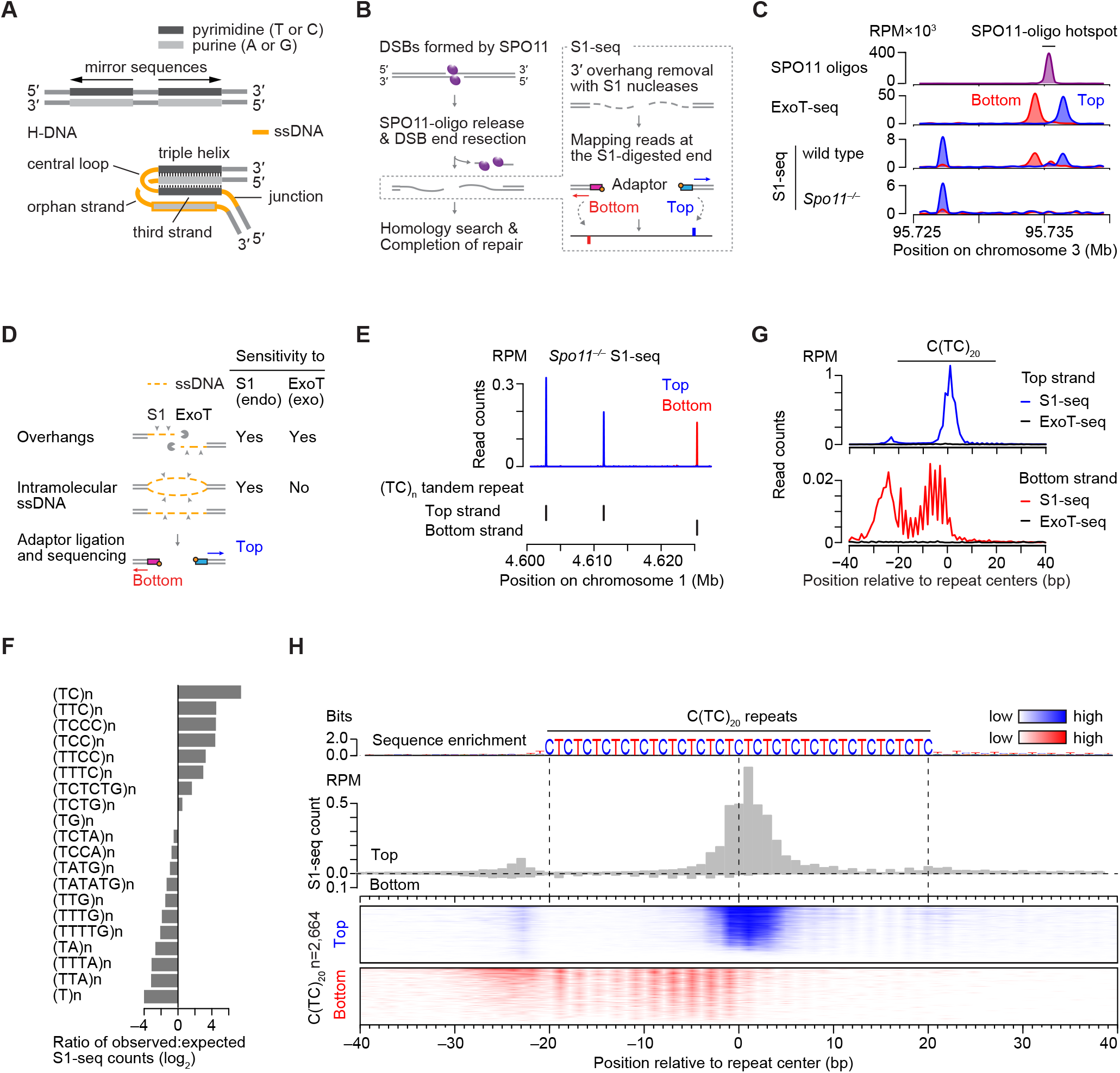
Spo11-independent S1-seq clusters at H-DNA motifs. **(A)** Schematic of H-DNA, consisting of a triplex and single-stranded DNA (ssDNA) regions, formed on a polypyrimidine/polypurine mirror sequence. **(B)** Early steps in meiotic recombination and overview of the S1-seq method. Spo11 (magenta ellipses) cuts DNA via a covalent protein-DNA intermediate. In S1-seq, sequencing adaptors are linked to duplex ends generated by removal of ssDNA tails using nuclease S1. Here and throughout, top strand refers to the strand that runs 5’ to 3’ in genomic coordinate space. The two adaptors shown are identical in sequence, but are color coded blue or red to indicate whether the resulting read maps to the top or bottom strand, respectively. **(C)** Strand-specific S1-seq (reads per million mapped reads [RPM]) at a representative DSB hotspot (right side, coincident with a peak in the SPO11-oligo sequencing) with a reproducible SPO11-independent read cluster nearby (left side). S1-seq and ExoT sequencing here and throughout are from Yamada et al. (2020). SPO11-oligo sequencing data here and throughout are from Lange et al. (2016). **(D)** Comparison of substrate specificities for nuclease S1 and ExoT. **(E)** Examples of pyrimidine-strand SPO11-independent S1-seq clusters at TC repeats. Black ticks below the plot show annotated TC repeats (RepeatMasker) on the top (+) or bottom (–) strand. **(F)** Preferential enrichment of S1-seq reads from *Spo11*^*−/−*^ mice at a subset of pyrimidine repeats (RepeatMasker annotations). **(G)** Averaged S1-seq and ExoT-seq signal around C(TC)_20_ sequences (n=2664). Note the different y-axis scales for top vs. bottom strand reads. **(H)** Stereotyped S1-seq read distributions at C(TC)_20_ sequences (n=2664). The sequence logo at the top indicates the base frequency in the sequences. The histogram (gray) shows the absolute average read count, illustrating the strong strand asymmetry. The heat maps below show the S1-seq reads separated by strand (pyrimidine strand in blue; purine strand in red). Each line is a single C(TC)_20_ sequence, ranked from highest total S1-seq read count at the top. The color gradients are calibrated separately for each strand to facilitate displaying the spatial patterning for the weaker S1-seq signal on the purine strand.

H-DNA motifs are enriched in the yeast and human genomes and are also common in many other species (Schroth and Ho, 1995; Clark et al., 2006). H-DNA motifs are overrepresented at introns and promoters (Bacolla et al., 2006; Lexa et al., 2014), coding regions of several disease-involved genes (Bissler, 2007), transposable elements (Kejnovsky et al., 2015), and fragile sites and oncogenic translocation breakpoints (Bacolla et al., 2016; Georgakopoulos-Soares et al., 2018). TC repeats, one type of H-DNA motif, are common in mammals (Tóth et al., 2000) and in 5’ noncoding regions of plant genomes (Zhang et al., 2006).

Intramolecular triplexes impinge on transcription, replication, and recombination (Mirkin and Frank-Kamenetskii, 1994; Frank-Kamenetskii and Mirkin, 1995). For example, (GAA)_n_ repeats stall replication in vivo (Krasilnikova and Mirkin, 2004) and cause transcriptional arrest (Sarkar and Brahmachari, 1992; Son et al., 2006). So-called “suicidal” mirror repeats can arrest DNA polymerase in vitro (Samadashwily et al., 1993). And H-DNA-forming sequences at the *c-MYC* locus interfere with transcription (Wang and Vasquez, 2004; Belotserkovskii et al., 2007).

H-DNA motifs are mutagenic (Wang and Vasquez, 2009). Triplex formation promotes genetic instability, mutation, and recombination leading to repeat expansion or genomic rearrangement (Bacolla and Wells, 2009). Transgenic mouse models indicate that H-DNA can induce genetic instability (Wang et al., 2008), while H-DNA motifs in the human genome are correlated with increased frequencies of somatic mutations, including recurrent mutations (Georgakopoulos-Soares *et al*., 2018).

H-DNA in eukaryotic genomes has been analyzed by various methods. Recognition of nuclei by triplex-specific monoclonal antibodies (Lee et al., 1987; Agazie et al., 1994) could be competed by exogenous triplex DNA (Agazie et al., 1996). Staining of interphase human cell nuclei with triplex-specific antibodies overlapped with sites of hybridization with probes intended to be specific for the displaced ssDNA (Ohno et al., 2002). And the dye Thiazole Orange, which binds to triplex DNA in vitro, also binds to dipteran chromosomal regions previously shown to be labelled by antibodies to triplex nucleic acid structures (Lubitz et al., 2010).

Unanswered questions include which motifs in the genome form H-DNA, and what are the specific triplex structures formed. A significant obstacle is the lack of methods to visualize H-DNA genome-wide at high resolution. Using potassium permanganate as a chemical probe for ssDNA, Kouzine et al. (2017) demonstrated that there is a propensity toward single-strandedness near H-DNA motifs in vivo. However, it is unclear whether the ssDNA detected was indeed H-DNA and, if so, what the structure of the H-DNA was, especially at individual loci.

In this study, we examined the triplex-forming potential of H-DNA motifs in mouse genomic DNA using S1-sequencing (S1-seq). In S1-seq, high-molecular-weight genomic DNA is embedded in agarose and digested with the ssDNA-specific nuclease S1, then adaptors are ligated to the blunted DSB ends for deep sequencing (Mimitou et al., 2017; Mimitou and Keeney, 2018; Yamada et al., 2020) (**Figure 1B**). We show here that S1-seq applied to DNA from mouse testis cells detects an unexpected but prominent signal at H-DNA motifs, and we use this signal to study structural features of intramolecular triplexes.

## Results

### SPO11-independent S1-seq signal

During meiosis, DNA double-strand breaks (DSBs) formed by the SPO11 protein are exonucleolytically processed to generate ssDNA tails that engage in homologous recombination (Lam and Keeney, 2014; Hunter, 2015; Cejka and Symington, 2021). We previously developed S1-seq to study this DSB resection (Mimitou *et al*., 2017; Mimitou and Keeney, 2018; Yamada *et al*., 2020). When applied to mouse testis samples, S1-seq reads were enriched near meiotic DSB hotspots in wild-type C57BL/6J (B6) mice relative to *Spo11*^*–/–*^ mice (**Figure 1C**) (Yamada *et al*., 2020). Because DNA ends on both sides of a DSB are resected, S1-seq signal mapping to the top strand spreads to the right from DSB hotspots, while bottom strand reads spread to the left.

However, we also observed S1-seq reads that were reproducibly enriched at sites distinct from meiotic DSB hotspots, that were similarly enriched in mice lacking SPO11, and that showed a strong strand bias (**Figures 1C and S1**). Little or no signal was seen at these sites if genomic DNA was digested with the 3’→5’ exonuclease ExoT (ExoT-seq) instead of the endonuclease S1, unlike around meiotic DSB hotspots (**Figures 1C and S1**). ExoT-seq signal is specific for DSBs (Canela et al., 2019), but the endonuclease S1 should be able to generate ligatable junctions at other ssDNA-containing structures in addition to DSBs, such as bubbles, nicks, or gaps (**Figure 1D**). Therefore, the lack of signal from ExoT-seq led us to infer that SPO11-independent S1-seq signal arises from a non-DSB structure(s) rather than from SPO11-independent DSBs such as those associated with replication or transposon activity.

In searching for features shared among DSB-independent S1-seq clusters (DISCs), we noticed that many were located at polypyrimidine repeats with the most prominent enrichment at TC repeats, and with a strong strand asymmetry in which most reads mapped to the pyrimidine strand (**Figure 1E**). When S1-seq read density at short tandem repeats in *Spo11*^*-/-*^ mice was calculated to evaluate this correlation more comprehensively, enrichment was observed for a subset of polypyrimidine repeats, including TC repeats (**Figure 1F**).

S1-seq showed a stereotyped signal distribution around TC repeats. To illustrate this, **Figures 1G and 1H** show the signal around all C(TC)_20_ repeats. (We chose C(TC)_20_ repeats for this example because there are many of them in the mouse genome and because they are mirror-symmetric.) On the pyrimidine strand, S1-seq showed a major cluster of reads at the repeat center, a minor cluster immediately to the left of the repeat, and a weak striated signal in the right half of the repeat. In contrast, the purine strand showed a weak, broad, striated enrichment within the left half of the repeat and a weak, diffuse signal in the sequence flanking the repeat on the left. We concluded that S1-seq likely detects intramolecular ssDNA preferentially formed in a sequence-dependent manner at pyrimidine-rich repeats. In principle, the ssDNA could have been present in vivo, but it is also possible that it mostly was formed ex vivo, when the genomic DNA was in agarose plugs in preparation for digestion with nuclease S1 (discussed further below).

### DISCs have sequence characteristics consistent with H-DNA

Many repetitive sequences contain motifs with propensity to form non-B-form structures that contain intramolecular ssDNA (**Figure 2A**) (Mirkin and Frank-Kamenetskii, 1994; Timsit and Moras, 1996; Herbert and Rich, 1999; Benham and Bi, 2004; Burge et al., 2006). We therefore speculated that DISCs may correspond to locations where non-B DNA forms readily. Based on the biochemical and biophysical properties of non-B DNA and in vitro experiments using plasmids and oligonucleotides, algorithms have been developed to identify sequences that have potential for non-B DNA isomerization (Kikin et al., 2006; Cer et al., 2011; Wang et al., 2013). When annotated non-B DNA motifs in the mouse genome (Kouzine et al., 2017) were compared with S1-seq signal for *Spo11*^*–/–*^ mice, intramolecular triplex (H-DNA) motifs harbored >50% of total S1-seq reads, much more than for other motifs (**Figure 2B**). Moreover, >99% of 144,478 DISCs (called with a cutoff of >10 hits per 50 bp) were located on putative H-DNA forming sequences (annotated H-DNA motifs plus other pyrimidine mirror repeats) (**Figure 2C**).

**Figure 2.**
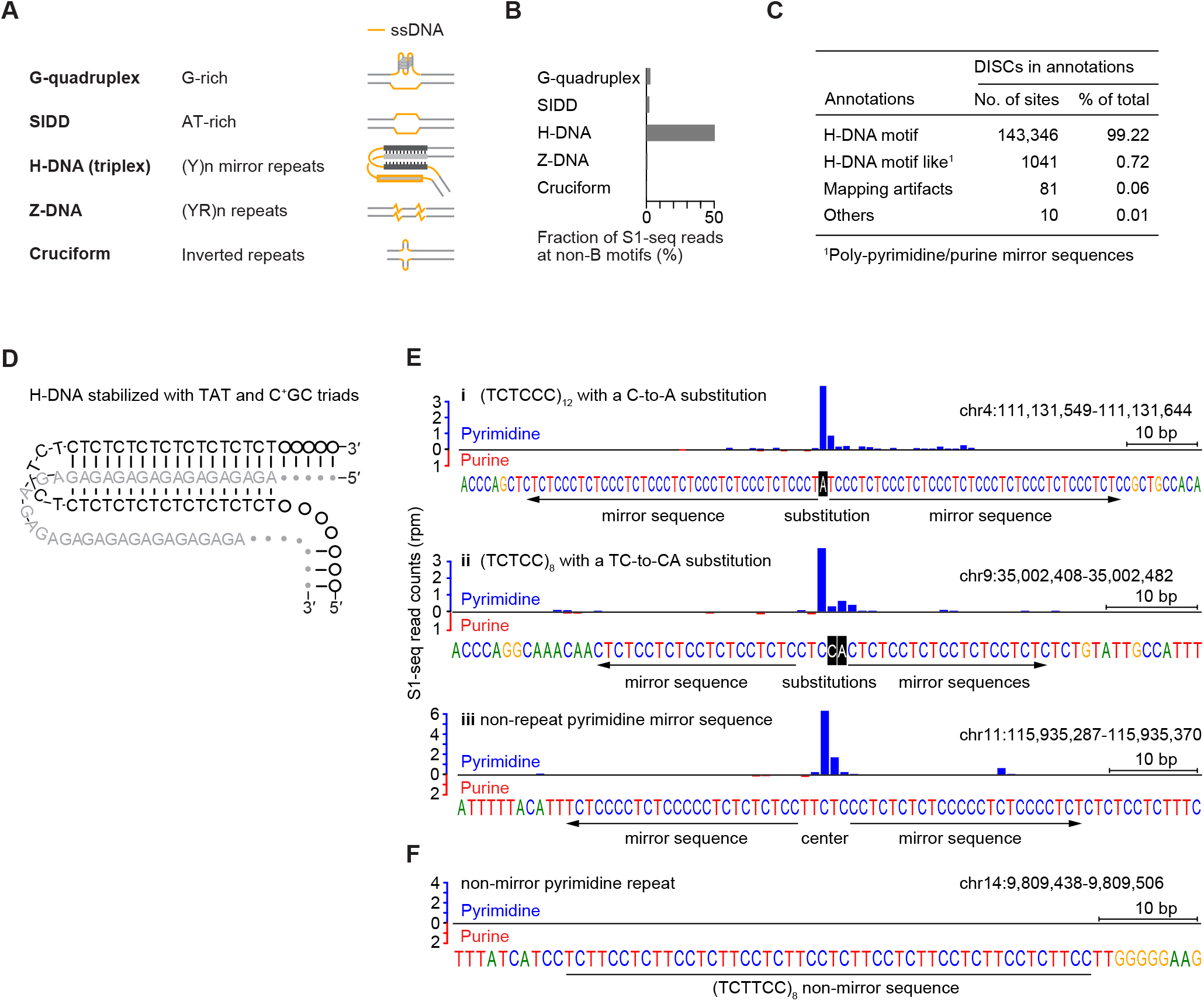
DISCs correspond to sites with characteristics consistent with H-DNA. **(A)** Schematic examples of non-B-form DNA structures containining ssDNA (yellow lines): G-quadruplex; stress-induced duplex destabilized site (SIDD); H-DNA; Z-DNA (left-handed double helix); cruciform. **(B)** Fraction of S1-seq reads from *Spo11*^*−/−*^ mice at non-B-form DNA motifs. Motif annotations are from (Kouzine *et al*., 2017). **(C)** Overlap of DISCs with H-DNA motifs. DISCs were called as peaks with >10 reads per 50 bp in *Spo11*^*−/−*^ S1-seq data, and compared with annotated H-DNA motifs (Kouzine *et al*., 2017). “H-DNA motif like” refers to polypyrimidine mirror repeats that were not annotated as H-DNA motifs. “Mapping artifacts” refers to DISCs with more randomly distributed S1-seq signal across long tandem repeats. “Others” are DISCs that could not be classified into these categories. **(D)** Schematic of H-DNA showing base sequence. H-DNA is stable if the constituent triads are TAT or C^+^GC in H-y isomers (shown), or are TAA or CGG in H-r isomers. As a result, H-DNA is readily formed on polypyrimidine mirror repeats. (**E**,**F**) Examples of S1-seq read patterns around different types of polypyrimidine sequences. (E.i and E.ii) Imperfect polypyrimidine repeats in which the interruptions to the repeat are centrally located. (E.iii) A mirror-symmetric polypyrimidine sequence that is not a direct repeat. (F) A polypyrimidine repeat that is not mirror symmetric.

Stable intramolecular triplex is favored by formation of Hoogsteen triads (**Figure S2A**), continuous runs of which can be formed by pyrimidine mirror sequences (**Figure 2D**) (Mirkin and Frank-Kamenetskii, 1994). Previous experiments showed that disrupting pyrimidine mirror repeats by base substitutions affected the H-DNA-forming potential of the repeats in plasmids, whereas substitutions located centrally in between mirror repeats did not (Mirkin et al., 1987). Consistent with these properties, we found that S1-seq reads were still enriched at polypyrimidine runs that had a centrally located substitution(s) that did not disrupt the mirror repeats (**Figure 2E.i and 2E.ii**). S1-seq reads were also enriched at non-repeat pyrimidine mirror sequences (**Figure 2E.iii**). However, non-mirror pyrimidine repeats showed little or no enrichment (**Figures 2F and S2B**). These results suggest that H-DNA-forming potential per se is predictive of S1-seq enrichment.

### S1-seq patterns may reflect properties of specific H-DNA isomers

We sought to discern how H-DNA structures might give rise to the observed S1-seq patterns, namely, the strand asymmetry and central position of the major cluster of reads at pyrimidine mirror sequences (**Figure 1H**). A pyrimidine mirror sequence can fold into four isomers (H-y5, H-y3, H-r5 and H-r3) named after the origin of the third strand in the triplex: whether it comes from the p<u>y</u>rimidine or pu<u>r</u>ine strand, and whether from the <u>5</u>’ or <u>3</u>’ half of that strand (**Figure 3A**) (Mirkin and Frank-Kamenetskii, 1994).

**Figure 3.**
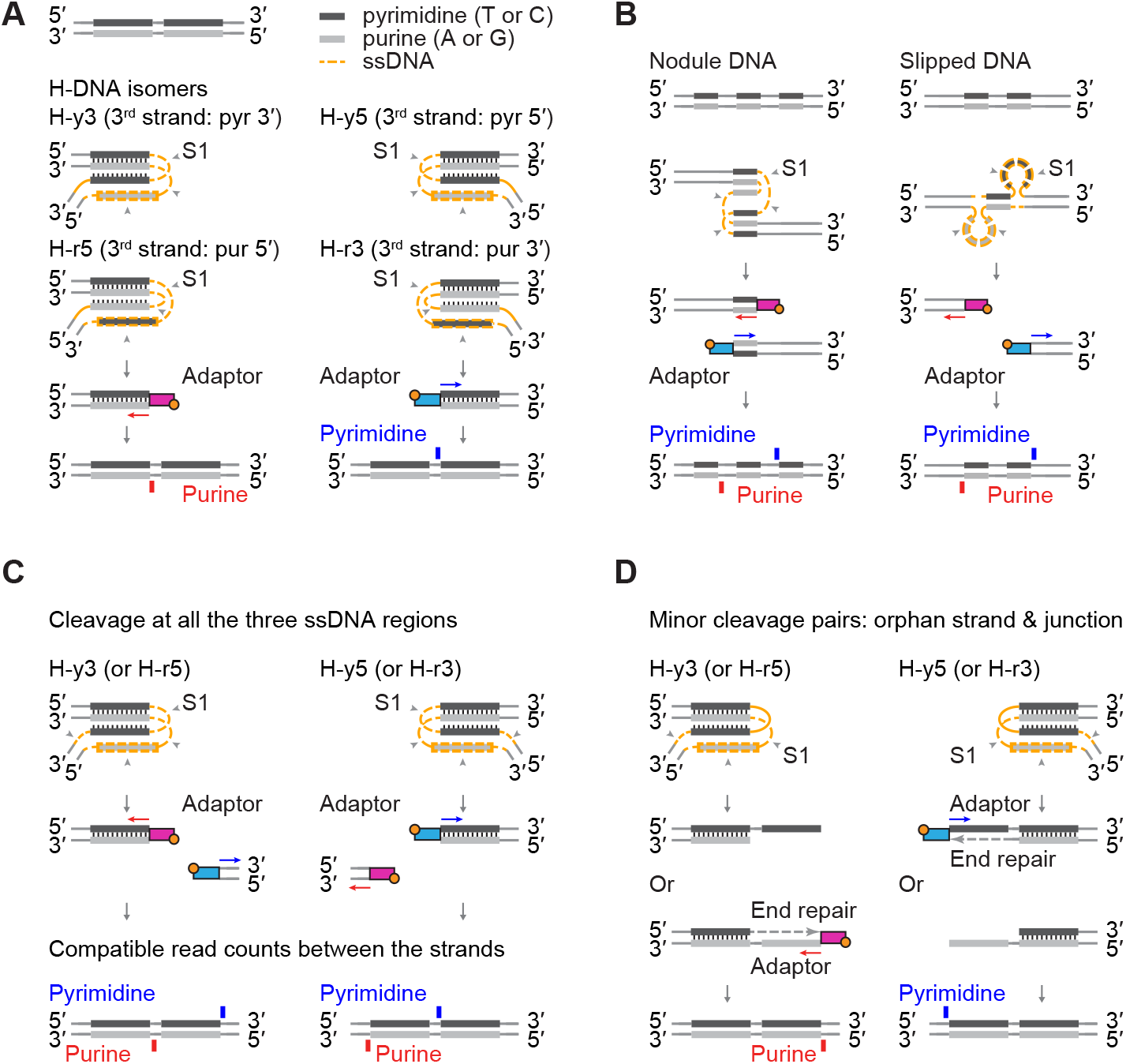
Models to illustrate how S1-seq strand asymmetry can be explained as a readout of H-DNA structure. In all panels, gray arrowheads indicate which ssDNA segments are digested with nuclease S1, and the adaptors are color coded to indicate whether the resulting sequencing read will map to the pyrimidine strand (blue) or purine strand (red). At the bottom of each panel, the schematic indicates the expected mapping position and strand for the S1-seq read. **(A)** H-DNA can exist as any of four isomers. Each isomer is named after the third strand in the duplex, e.g., if the third strand is from the 5’ half of the pyrimidine strand, the isomer is called H-y5. For H-y5 and H-r3 (right side), S1 digestion to yield a duplex DNA end at the border between the central loop and the original triplex should allow adaptor ligation to the right half of the mirror sequence, resulting in a centrally located pyrimidine-strand S1-seq read (blue). In contrast, digestion of isomers H-y3 and H-r5 should allow adaptor ligation to the left half of the mirror sequence, resulting in a central purine-strand read (red). **(B)** Other possible non-B-form structures, nodule DNA and slipped DNA, were not suitable for explaining the position and strand asymmetry of S1-seq reads. **(C)** Expected S1-seq read positions if ssDNA of H-DNA is fully digested. For H-y5 and H-r3 (right side), digestion of both strands at the junction should yield a flanking purine-strand read in addition to the central pyrimidine-strand read detailed in panel A. For H-y3 and H-r5 (left side), the junction read should be map to the pyrimidine-strand. If junction digestion is efficient, then comparable read counts are expected at the junction and central positions. **(D)** Expected S1-seq read positions if H-DNA is only partially digested by nuclease S1 on the orphan strand and junction. Partial digestion at the indicated positions on H-y5 (right) would yield a 5’ overhang that could be filled in by T4 DNA polymerase and ligated to a sequencing adaptor, yielding an S1-seq read that maps to the pyrimidine strand just to the left of the mirror repeat sequence. For H-y3, this scenario predicts a purine-strand read to the right of the mirror repeat, while H-r5 and H-r3 would yield 3’ overhangs. If these are inefficiently polished, they would not yield a sequencing read (as shown); if polished and ligated, they would yield a central read indistinguishable from S1 having cleaved the central loop (**Figure 3A**).

Complete digestion of central loop ssDNA regions with nuclease S1 and dissociation of the triplex should leave (among other things discussed below) a duplex DNA end at the border between the central loop and the original triplex (**Figure 3A**). For isomers H-y5 and H-r3, this duplex DNA end should allow adaptor ligation to the right half of the mirror sequence, resulting in a centrally located pyrimidine-strand S1-seq read (**Figure 3A right**; blue arrow). This matches the observed pattern in S1-seq data (**Figure 1H**), thus, H-y5 and H-r3 can plausibly account for the major central pyrimidine-strand S1-seq signal. In contrast, digestion of isomers H-y3 and H-r5 should allow adaptor ligation to the left half of the mirror sequence, resulting in a central purine-strand read (**Figure 3A left**, red arrow). This was not observed at high frequency, so these isomers are unlikely to be sources of S1-seq signal.

Polypyrimidine mirror sequences can also make non-B-form structures other than H-DNA, namely nodule DNA and slipped DNA, which are tandem repeats of triplexes or of ssDNA loops, respectively (**Figure 3B**) (Panyutin and Wells, 1992; Sinden et al., 2007). These structures are symmetric, unlike H-DNA, so they would be expected to produce equivalent numbers of S1-seq reads from both strands. Moreover, such reads would not be centrally located within the mirror sequence (**Figure 3B**). Therefore, neither of these structures is a good candidate to explain the central S1-seq signal.

DISCs show strong asymmetry between the strands for total S1-seq read count in and around pyrimidine mirror sequences (**Figure 1H**). If S1 cleaves both strands of the H-DNA junction region, it should leave a duplex DNA end that flanks the mirror sequence and that can be ligated to an adaptor, giving rise to a sequencing read pointed away from the mirror sequence (**Figure 3C**). This would yield a purine-strand read on the left side for H-y5 and H-r3 (the inferred major source of DISC signal) or a pyrimidine-strand read on the right for H-y3 and H-r5 (**Figure 3C**). Such reads consistent with H-y5 and H-r3 were indeed observed (**Figure 1H**). However, if S1 cleavage of junctions is efficient, these reads should be comparable in number to the reads from cleavage at the central loop, but we instead observed a much smaller number of these candidate junction reads (**Figures 1E, 1G, and 1H**).

A straightforward way to account for this difference is if S1 does not always fully digest all of the ssDNA segments. For example, if S1 cleaves the loop and orphan strand but not the junction strand, this would leave a long 3’ overhang for H-y5 that might block ligation if it is inefficiently removed during the subsequent polishing step with T4 DNA polymerase (**Figure S3A**). If the junction strand is less efficiently digested than the other strands, this scenario would represent a majority of S1-digested H-DNA molecules, which might then account for the relative deficit of junction reads as compared to central reads. Additionally, if H-DNA were sometimes digested on the orphan and junction strands but leaving the central loop intact, this would be expected to leave a long 5’ overhang for H-y5 that, after fill in by T4 DNA polymerase, would be predicted to yield a pyrimidine-strand junction read mapping just to the left of the mirror repeats (**Figure 3D**). Pyrimidine-strand reads matching this expectation are apparent on the left side of C(TC)_20_ repeats (**Figures 1G and 1H**). Depending on which strands end up being cleaved by S1 and where those cleavages occur, additional detailed features of the DISC S1-seq patterns can be readily explained, including the periodically spaced purine-strand reads inside the left half of mirror sequences (**Figure S3B**).

To further test whether DISCs might reflect specific H-DNA structures, we compared the location and strength of the S1-seq signal with the previously determined sensitivity of an H-y5-forming plasmid to chemical probes KMnO_4_, which detects unpaired T and C, and acid depurination, which detects unpaired G and A. Glover et al. studied the chemical reactivity of the H-y5 isomer formed on (TC)_17_ (Glover et al., 1990). On the pyrimidine strand, chemical reactivity was highest in the central loop region, followed by flanking sequence to the left, then within the 5’ (left) half of the repeat (**Figure S3C**). On the purine strand, chemical reactivity was highest in the central loop region and the orphan strand (the 3’ (left) half of the repeat), then the 5’ (right) half of the repeat, then downstream to the right. The greater reactivity of the central loop than the junction sequence is consistent with the hypothesis that the central loop might also be more easily digested by nuclease S1. The modest chemical reactivity of both the purine strand and the third strand in the triplex suggests that, even though fully incorporated in the triplex in the H-DNA structure model, they may be partially unpaired in solution and thus may also be able to be cleaved by S1 nuclease. Taken together, the S1-seq results appear to be largely congruent with expectation based on chemical probes of single-strandedness in H-y5 H-DNA.

Nevertheless, we noticed rare examples that might represent (mixtures of) other H-DNA isomers. Loop sequence plays a crucial role for isomerization of intramolecular DNA triplexes in supercoiled plasmids (Kang and Wells, 1992; Shimizu et al., 1994). **Figure S4A** shows an example of a pyrimidine mirror repeat with a 10-bp interruption between the repeats. The S1-seq signal is markedly different from the pattern at more contiguous pyrimidine mirror repeats, with central signal enriched on both strands. This pattern could be consistent with alternative H-DNA isomers (i.e., H-y3 or H-r5 in addition to the more prevalent H-y5 or H-r3).

### S1-seq patterns at pyrimidine mirror repeats with different sequence compositions

H-DNA studies began with the discovery that some plasmids showed reactivity to S1 nuclease (Weintraub, 1983). Subsequently, H-DNA structural features have been extensively studied using plasmids in which various potential H-DNA-forming sequences were individually cloned and characterized (Wells et al., 1988). However, the number of sequences analyzed per study was of necessity limited, and whether results with a given set of sequences could be extrapolated to untested sequences remains unclear. S1-seq of mouse genomic DNA allows us to systematically analyze many thousands of pyrimidine mirror repeats over the mouse genome simultaneously.

First, we consider the effect of repeat sequence on S1-seq signal. Because S1-seq read counts varied between different pyrimidine mirror sequences (**Figure S2C**), we hypothesized that S1-seq signal reflects the frequency and stability of triplex structure formation. To test this, we compared the S1-seq signal intensity with the ease of H-DNA formation in plasmids. Plasmids with (TCCTC)_n_ require greater superhelical stress per unit length to form H-DNA than with the repeat (TC)_n_ (Collier and Wells, 1990). Congruently, we found that the S1-seq signal intensity was greater for TC repeats than for TCCTC repeats of the same length (compare C(TC)_24_ with TC(TCCTC)_9_T in **Figure S2B**).

Moreover, detailed spatial patterns of S1-seq reads differed between different repeat sequences. Specifically, each type of repeat showed a periodically spaced S1-seq signal that was highly stereotyped for a given repeat sequence, but that differed between repeats of different sequence (**Figures 1H and S4B**). Analogously, Collier and Wells (1990) found that plasmid-borne (TCCTC)_12_ gave a periodic pattern of chemical sensitivity with peaks every five bases. A simple single-base preference for the chemical probes or S1 nuclease alone would not explain why different repeat sequences give peaks at different position, so we surmise that local differences in triplex structure modulate nuclease sensitivity and chemical reactivity.

### Variations in S1-seq patterns according to pyrimidine mirror repeat length

Next, we consider the effect of repeat length on S1-seq signal. How the lengths of mirror repeats shape H-DNA three-dimensional structures remains largely uncharacterized. Previous plasmid studies suggested that long polypyrimidine stretches may form large H-DNA (Han and de Lanerolle, 2008) or nodule DNA (two tandem H-DNAs) (Panyutin and Wells, 1992). To address this issue, we focused on TC repeats of various lengths because these repeats are abundant in the mouse genome and are highly enriched for S1-seq reads.

**Figure 4A** shows a heat map of strand-specific S1-seq reads for C(TC)_n_ repeats with n ranging from 16 to 38 in steps of 2, centered on the repeat midpoints. The read distributions were strongly stereotyped across copies of the same length, but showed progressive differences that tracked with repeat length. Specifically, the strong cluster of pyrimidine-strand reads inferred to be from cleavage at the central loop grew wider as TC repeat length increased, and the weaker clusters of pyrimidine- and purine-strand reads inferred to represent junction cleavage moved progressively leftward.

**Figure 4.**
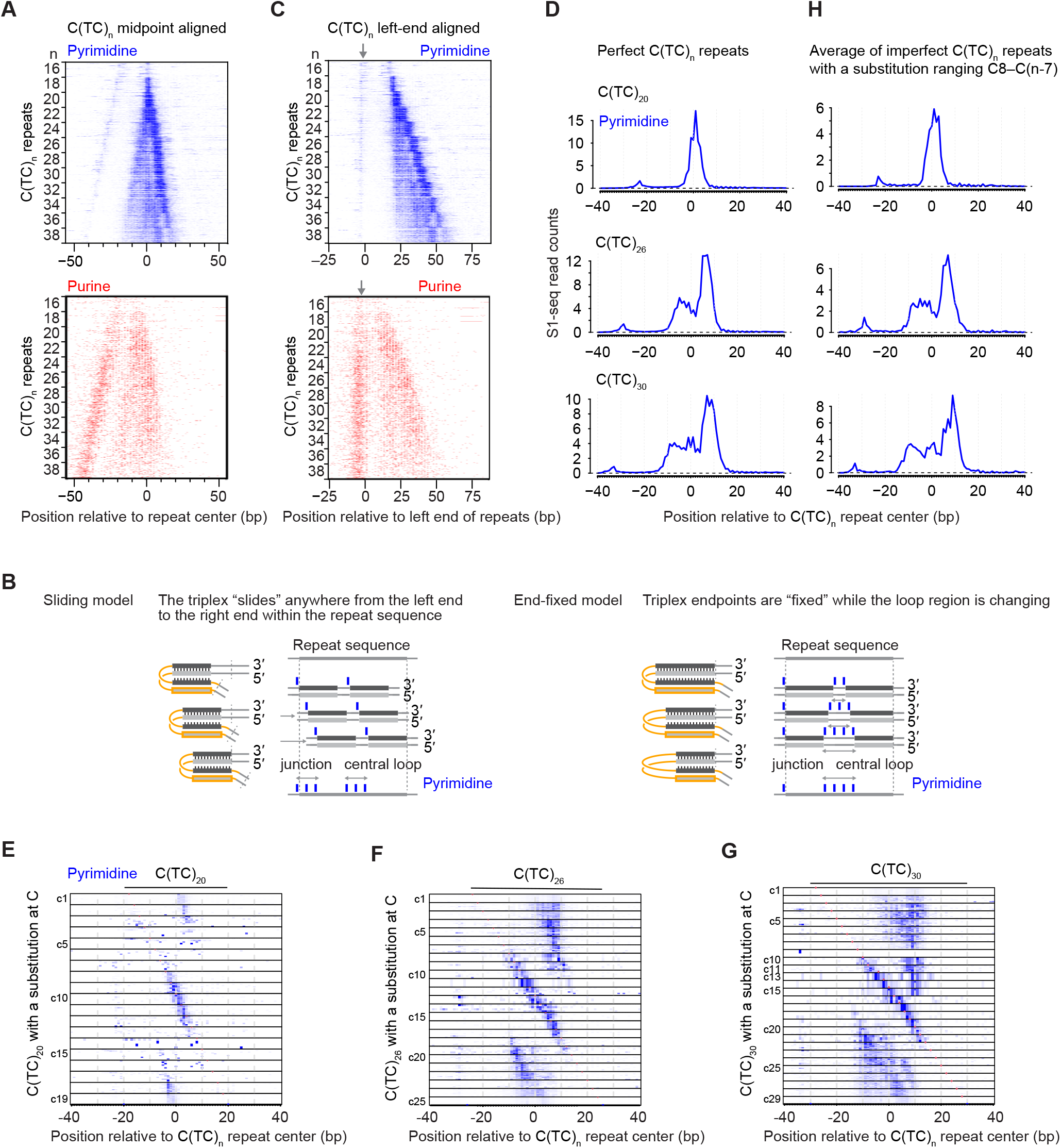
Variation in S1-seq read patterns with differences in TC repeat length and composition. **(A)** Strand-specific heat maps of S1-seq signal for C(TC)_n_ repeats with n ranging from 16 to 38 in steps of 2, centered on the repeat midpoints. Color gradients for pyrimidine the strand (blue) and purine strand (red) are scaled differently to illustrate each strand’s spatial patterns. For each value of n, 100 loci of that length were randomly selected for display. **(B)** Alternative models to explain the spread of pyrimidine-strand central signal as TC repeat length increases. See text for details. The triplex length is shown as constant for simplicity, but it can vary in either model. **(C)** Fixed position of junction reads on both strands irrespective of TC repeat length. The heat maps show strand-specific S1-seq signal for the same C(TC)_n_ repeats as in panel A, but lined up along the start of the repeat. Gray arrows indicate the position of the presumed junction reads. **(D)** Averaged pyrimidine-strand S1-seq signal (read counts) for perfect C(TC)_n_ repeats of the indicated lengths. (**E, F, G**) S1-seq patterns on imperfect C(TC)_n_ repeats differ in stereotyped ways depending on the position of the imperfection and the length of the repeat. Heat maps show pyrimidine-strand S1-seq maps for imperfect C(TC)_20_ (panel E), C(TC)_26_ (panel F), and C(TC)_30_ (panel G). Each repeat analyzed has a single cytosine substitution, and the substitutions positions are grouped and ordered with the leftmost substation (C1) at the top. (**H**) Averaged pyrimidine-strand S1-seq signal (RPM) for imperfect C(TC)_n_ repeats of the indicated lengths. For each graph, repeats with a single cytosine substitution ranging from C8 to C(n-7) were averaged.

We considered two models to explain the expansion of the central S1-seq signal, taking into account that C(TC)_16_ is the minimum needed to form detectable H-DNA (Htun and Dahlberg, 1989). In one model (the “sliding” model, shown in **Figure 4B left**), the triplex segment can be positioned anywhere within a repeat that is longer than the minimum (i.e., n ≥ 16). As a result, the central loop can occupy any position within the repeat except within n = 8 (16 bp) from the ends, and the central S1-seq signal becomes wider at longer repeats because it is the superposition of a population of alternative H-DNA structures that can be differentiated from each other by “sliding” the triplex across the repeat. An alternative model (the “end-fixed” model, shown in **Figure 4B right**) envisions that the triplex position is fixed at the left and right ends of the repeat. In this model, the longer the repeat is, the longer the central loop will be on average and the wider the distribution of central loop boundary positions will be. This model thus explains the spread of the central S1-seq signal as the superposition of a population of H-DNA structures that have the triplex position fixed at the ends, but variable positions for the boundary of the central loop. In both models, the triple-stranded region can vary in length.

To determine which model is better suited to explain the overall signal pattern, we focused on the clusters of junction reads immediately to the left of the repeat on both strands. The sliding model predicts that the junction position will be variable as the repeat length increases, spreading from the left end of the repeat into the repeat itself. In contrast, the end-fixed model explicitly posits a uniform junction position just outside the left end of the repeat, regardless of the length of the repeat. When the heatmaps were centered on the left end of the polypyrimidine tracts, the junction S1-seq signal remained confined to one position irrespective of the number of repeats (**Figure 4C**), consistent with the end-fixed model.

### Various intramolecular triplex structures form on long TC repeats

For longer TC repeats, the central S1-seq signal on the pyrimidine strand showed pronounced substructure (**Figure 4A**), with multiple peaks when repeats of the same length were averaged (**Figure 4D**). This substructure cannot be explained by a simple end-fixed model in which the triplex length is also fixed and the central loop region is entirely ssDNA. We therefore hypothesized instead that the complex S1-seq read distribution reflects an average over a population of alternative structures that differ with respect to which portions of the central loop are single stranded.

To test this idea, we examined the effects of repeat-disrupting substitutions. As noted above, substitution is tolerated within the central loop (**Figure 2E**), but a substitution within the triplex-forming region disfavors H-DNA formation because the substitution prevents Hoogsteen triad formation (Mirkin *et al*., 1987; Belotserkovskii et al., 1990). We reasoned that a repeat-disrupting substitution would limit the variety of alternative structures that might be formed by a long TC repeat because the triple-stranded region would be constrained not to overlap the substitution. We therefore searched for imperfect C(TC)_n_ repeats in which one cytosine position was instead a different base, and then examined how S1-seq read patterns changed depending on the position of the substitution.

Heatmaps for imperfect C(TC)_n_ repeats are shown in **Figures 4E, 4F, and 4G** for n = 20, 26, and 30, respectively. The leftmost C in C(TC)_n_ is indicated as C0, and each cytosine is named from C0 to Cn in order from left to right. As predicted, each substitution position was associated with a marked and stereotyped difference in the S1-seq read distribution. Substitutions near the center of the repeat (e.g., C10 to C21 for C(TC)_30_) gave pronounced clusters of pyrimidine-strand S1-seq reads immediately around the substitution, as expected if the substitution is constrained to be within the central loop and is often single-stranded.

If the complex pattern for the central S1-seq signal on long perfect TC repeats reflects a population of alternative structures with ssDNA occupying different positions, we reasoned that we could mimic this more complex population by combining the simpler populations observed for the imperfect TC repeats whose central loops are more constrained. Indeed, when we created average plots combining imperfect repeats with central substitutions, these recapitulated well the average plots for perfect repeats for the pyrimidine strand (**Figure 4H**) and the purine strand (**Figures S5A, S5B**).

### S1-seq patterns at H-DNA motifs are similar in resting vs. activated B cells

To ask whether DISCs might reflect H-DNA that was already present in vivo, we measured S1-seq patterns around H-DNA motifs in DNA isolated from cultured primary mouse splenic CD43+ B cells, comparing transcriptionally quiescent resting cells to transcriptionally active cells that had been stimulated by treatment ex vivo with lipopolysaccharide and interleukin 4 (LPS+IL4) (Kouzine et al., 2013). Previous studies using ssDNA-seq—which uses KMnO_4_ and nuclease S1 treatment to introduce DSBs at locations enriched for unpaired bases in vivo—detected a broad zone of sequencing read enrichment around H-DNA motifs in activated B cells, compared with a narrower band of relative depletion in resting cells (**Figure 5A**) (Kouzine *et al*., 2017). We therefore reasoned that S1-seq patterns would also be different between the two cell states if S1-seq were detecting transcription-promoted H-DNA that had formed in vivo. However, average S1-seq signal intensity at H-DNA motfis was highly similar between activated and resting B cells (**Figure 5A**), and spatial patterns and read density were also similar when considering just C(TC)_20_ repeats (**Figures 5B, 5C, and 5D**). These data thus did not provide support for the idea that S1-seq captures intramolecular triplex structures that were formed in vivo. One possibility is that most or all of the triplexes detected by S1-seq form after DNA isolation. Alternatively, H-DNA detected by S1-seq may form in vivo, but independently of transcriptional status. Importanly, these findings speak only to what S1-seq detects; they do not exclude that H-DNA does indeed form in vivo.

**Figure 5.**
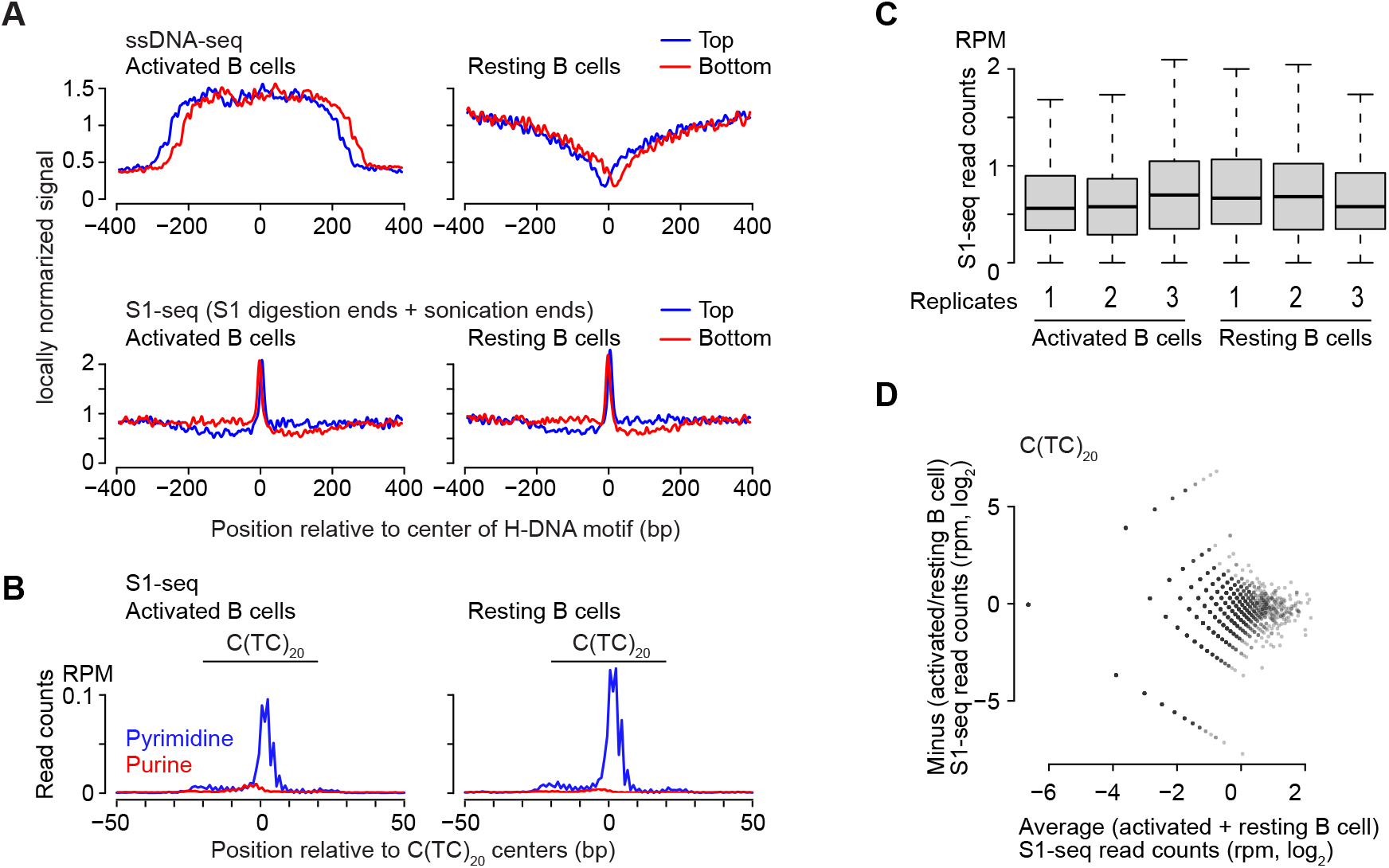
S1-seq patterns at H-DNA motifs in activated vs. resting splenic B cells. **(A)** Locally normalized average plots of ssDNA-seq (Kouzine *et al*., 2017) and S1-seq for activated and resting B cells, centered on H-DNA motifs as annotated by Kouzine et al. (2017). **(B)** Average plot of S1-seq for activated and resting B cells, centered on C(TC)20 loci. **(C)** Similar S1-seq read counts at C(TC)_20_ loci in activated vs. resting B cells. Box plots are as defined in the legend to **Figure S2B**. Three biological replicates are shown for each cell state. **(D)** MA plot of S1-seq read counts at C(TC)_20_ loci comparing activated and resting B cells.

If the H-DNA detected by S1-seq was formed ex vivo, a couple of observations suggest that having a pyrimidine mirror sequence may not be sufficient to yield detectable signal. First, different TC repeats of the same length yielded widely different amounts of S1-seq signal (**Figure S2B**). Second, the single TC repeat longer than C(TC)_20_ in the yeast genome did not yield characteristic H-DNA S1-seq patterns (data not shown).

### Phylogenetic aspects of TC repeats

Noticing that mice have many more TC repeats than humans prompted us to investigate the phylogenetic distribution of TC repeats. C(TC)_20_ is abundant (>2500 copies) in the genome assemblies of mouse, rat, and Chinese hamster, but is much less frequent in more distantly related species including kangaroo rat, naked mole-rat, ground squirrel, and pika (**Figure 6A**).

**Figure 6.**
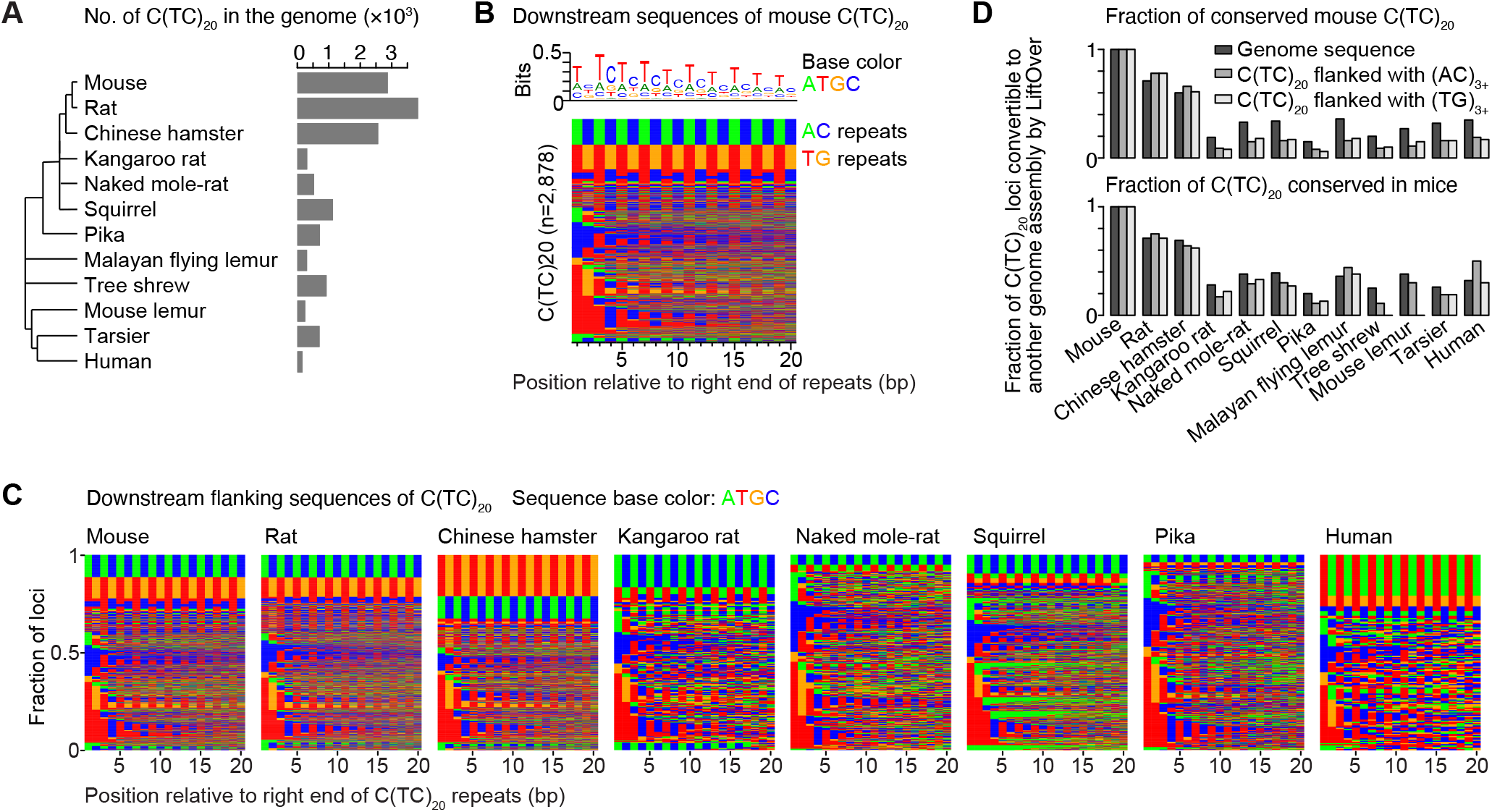
Amplification of TC repeat copy number in rodents. **(A)** C(TC)_20_ repeat copy number in various species: mouse (C57BL/6J strain of *Mus musculus*), rat (*Rattus norvegicus*), Chinese hamster (*Cricetulus griseus*), kangaroo rat (*Dipodomys ordii*), naked mole-rat (*Heterocephalus glaber*), squirrel (*Spermophilus tridecemlineatus*), pika (*Ochotona princeps*), Malayan flying lemur (*Galeopterus variegatus*), tree shrew (*Tupaia belangeri*), mouse lemur (*Microcebus murinus*), tarsier (*Tarsius syrichta*), and human (*Homo sapiens*). Tree topology is from the UCSC genome browser; branch lengths are not to scale. **(B)** Sequence context of mouse C(TC)_20_ repeats. The clustered color map shows the 20 nucleotides 3’ of C(TC)_20_ repeats in mice. Green, blue, red, and orange indicates adenine, cytosine, thymine, and guanine, respectively. **(C)** Sequence context of C(TC)_20_ repeats in other species. The clustered color maps are presented as in panel B. **(D)** Conservation of C(TC)_20_ sequences. The top graph shows the fraction of mouse C(TC)_20_ sequences (followed by TG or AC repeats, as indicated) that are conserved in each of the indicated species. The bottom graph shows the fraction of such C(TC)_20_ sequences from the indicated species that are conserved in mouse. Black bars show overall genomic sequence conservation as a point of comparison.

For many mouse C(TC)_20_ repeats, the sequence immediately downstream (3’ on the pyrimidine strand) is enriched for additional degenerate TC repeats, but a substantial subset (648 out of 2,878 total C(TC)_20_ repeats) instead has an AC repeat in either forward or reverse orientation (i.e., either a TG or AC repeat) (**Figure 6B**). The same is true for rat and Chinese hamster (**Figure 6C**). Moreover, more than 60% of mouse C(TC)_20_ repeats that are followed by TG or AC repeats are conserved in rat and Chinese hamster (**Figure 6D, top**), and, conversely, more than 60% of such repeats in rat and Chinese hamster are conserved in mouse (**Figure 6D, bottom**). Some C(TC)_20_ repeats in kangaroo rat, naked mole-rat, and squirrel were also followed by AC repeats, but downstream TG repeats were rare (**Figure 6C**), and C(TC)_20_ repeat positions were less well conserved with mice (**Figure 6D**). In the human genome, C(TC)_20_ repeats were often followed by AT or, less often, TG repeats (**Figure 6C**). These findings indicate that TC repeats amplified and spread throughout the genome at or before the last common ancestor of mice, rats, and Chinese hamster. Moreover, at least a subset of TC repeats were in the context of a more complex repeat structure at the time of amplification.

## Discussion

In this study, we uncovered features of apparent triplex structures within the mouse genome by using S1-seq signal as a footprint of non-B-form DNA structure. DISCs—S1-seq hotspots that appear to be unrelated to the presence of DSBs—were clearly correlated with the H-DNA forming potential of their DNA sequences. Moreover, local S1-seq distributions were consistent with H-y5 and/or H-r3 isomers of H-DNA, including the strong central pyrimidine-strand signal and the strong strand asymmetry of the sequencing read count.

An open question is whether the triplex detected by S1-seq was already present in vivo, or if instead it formed ex vivo. A plausible argument can be made that most, if not all, of the detected triplexes were formed ex vivo, because the removal of histones and other proteins from the chromatin while the DNA was constrained in agarose might have provided sufficient negative superhelicity to favor triplex formation. Moreover, the digestion with nuclease S1 was conducted at acidic pH in the presence of divalent cations, which both favor H-DNA formation (discussed further below). We sought evidence in support of triplex formation in vivo by comparing S1-seq patterns on DNA from resting vs. activated B cells. The lack of a clear difference between the cell types indicates either that most of the DISC signal was formed ex vivo, or that H-DNA formation in vivo is independent of transcriptional status.

Taken together, we favor the idea that most of H-DNA detected by S1-seq was formed ex vivo. Importantly, however, we cannot exclude the possibility that at least some triplex was already present in vivo. Moreover, even if all of the triplex detected by S1-seq had formed ex vivo, it would not exclude the possibility that triplex does indeed form in vivo. Nevertheless, our findings establish that S1-seq can be useful as a new tool to probe H-DNA potential genome-wide at high resolution.

The S1-seq signal is consistent with both H-y5 and H-r3 isomers, but the following considerations lead us to infer that the triplex detected by S1-seq is probably H-y5. For formation of H-y isomers, acidic pH and longer length are favorable (Collier and Wells, 1990; Mirkin and Frank-Kamenetskii, 1994). Moreover, the H-y5 isomer is observed at lower superhelical density than H-y3, and can even occur on linearized plasmids (Htun and Dahlberg, 1988). Also, the presence of divalent cation (e.g., Ca^2+^, Zn^2+^, Mg^2+^, and Mn^2+^) makes the H-y5 isomer preferable (Kang et al., 1992). Relevant to these points, the nuclease S1 digestion buffer is acidic (pH 4.5) and contains 4.5 mM Zn^2+^.

In contrast, H-r isomers are stable at neutral pH in the presence of divalent metal cations, with H-r3 requiring a higher super helical density than H-r5 (Htun and Dahlberg, 1989). Importantly, CG*A^+^ base triads can form H-r isomers at acidic pH without the presence of cation (Beal and Dervan, 1991; Dayn et al., 1992; Malkov et al., 1993; Klysik, 1995). As a result, (TA)_10_(TC)_10_ can form the H-r3 isomer and (TC)_10_(TA)_10_ can form the H-r5 isomer. However, S1-seq reads were not enriched at either of these sequences (data not shown). This lack of S1-seq signal suggests either that H-r3 does not form, or that it is not detected by S1-seq when it is present. By process of elimination, H-y5 appears to be the more likely source of S1-seq signal.

We note that the triplex-forming sequences detected by S1-seq in this study are longer than most of the sequences that have been examined in plasmids in previous studies. For TC repeats longer than (TC)_15_, one negative superhelical turn in plasmid DNA is relieved for every 11 nucleotides of (TC)_n_ repeat that can be converted from duplex to the H-DNA conformation (Htun and Dahlberg, 1989). TC repeats shorter than (TC)_13_ relieve more superhelical turns to form H-DNA, suggesting that the longer repeats (longer than (TC)_15_) can form H-DNA at lower superhelical density. We found that S1-seq signal was enriched at TC repeats longer than C(TC)_16_, consistent with triplex formation that requires only a low superhelical density. The H-y5 structure is favored under conditions of very low or no topological stress (explained by predicted increased stacking energy in the H-y5 conformer), whereas high topological stress generated only the H-y3 conformer (which relaxes more supercoil writhe) (Broitman, 1995). These considerations lend further support to the conclusion that S1-seq is detecting primarily the H-y5 conformation, and also reinforces the plausibility of H-DNA forming on the agarose-embedded linear genomic DNA under the conditions of S1 digestion..

S1-seq (this study) and ssDNA-seq (Kouzine *et al*., 2017) both detect ssDNA but gave different results at H-DNA motifs. First, S1-seq signal was enriched at most of the long H-DNA motifs, but ssDNA-seq was enriched at only a subset of the motifs and showed only a weak enrichment at TC repeats. Second, ssDNA-seq showed marked differences between resting and activated B cells, but S1-seq did not. Third, DSB-independent S1-seq signal was almost exclusively at H-DNA motifs, but ssDNA-seq signal was observed at H-DNA as well as other non-B motifs (Kouzine *et al*., 2017). These differences presumably reflect the different methodologies for probing ssDNA. In particular, ssDNA-seq using nuclease S1 to detect sites where ssDNA was previously modified in vivo with permanganate, whereas S1-seq relies on direct digestion of unmodified DNA with nuclease S1.

Our findings show that S1-seq can be used to study detailed triplex structures on mammalian genomic DNA. One strength is that S1-seq permits examination of a large number of intramolecular-triplex forming sequences on genomic DNA at once. Thus, this study may serve as a basis for investigating the function of triplex structures on genomic DNA.

## Methods

### Mice

Experiments conformed to the US Office of Laboratory Animal Welfare regulatory standards and were approved by the Memorial Sloan Kettering Cancer Center Institutional Animal Care and Use Committee. Mice were maintained on regular rodent chow with continuous access to food and water until euthanasia by CO_2_ asphyxiation prior to tissue harvest. C57BL/6J (stock no. 000664) male mice were obtained from the Jackson Laboratory.

### S1-seq

#### B cell preparation

Splenic B cells were obtained as described previously (Kouzine *et al*., 2013). Briefly, resting naive mouse B cells were isolated from splenocytes with anti-CD43 MicroBeads (Miltenyi Biotec) by negative selection and activated for 72 hr in the presence of LPS (50 μg/ml final concentration, E. coli 0111:B4; Sigma-Aldrich) plus IL4 (2.5 ng/ml final concentration, Sigma-Aldrich). Apoptotic cells were removed with a Dead Cell Removal Kit (Miltenyi) followed by Ficoll gradient with >90% live-cell purity.

#### DNA extraction in plugs

Cells were embedded to protect DNA from shearing, and DNA was liberated by treatment with SDS and proteinase K as described previously (Yamada *et al*., 2020). In-plug overhang removal with nuclease S1 and adaptor ligation were performed as described previously (Mimitou *et al*., 2017; Mimitou and Keeney, 2018). After ligation to biotinylated adaptros, DNA was purified from the agarose, sheared by sonication, purified with streptavidin, ligated to second-end adaptors, amplified, and sequenced as described previously (Mimitou *et al*., 2017; Mimitou and Keeney, 2018; Yamada *et al*., 2020).

#### Mapping and preprocessing

Sequence reads were mapped onto the mouse reference genome (mm10) by bowtie2 version 2.2.1 (Langmead et al., 2009) with the argument –X 1000. Uniquely and properly mapped reads were counted, at which a nucleotide next to biotinylated adaptor DNA was mapped.

### Sequence motif search

Genome sequences were searched using dreg (EMBOSS version 6.6.0.0) (Rice et al., 2000) with the default parameters. The repeat length was defined as that of the longest repeating subsequences with no mismatches, insertion or deletions, i.e. no (TC)_20_ repeats were annotated within a (TC)_21_ repeat. Genome versions used in this study were listed in Supplementary Table S1.

### Other data sets and data availability

Raw and processed sequencing S1-seq data were deposited at the Gene Expression Omnibus (GEO) (accession number is pending). We used SPO11-oligo, yeast S1-seq data from GEO accession numbers GSE84689 and GSE85253, respectively; and ExoT-seq and mouse S1-seq data from GSE141850 (Lange et al., 2016; Mimitou *et al*., 2017; Yamada *et al*., 2020). Non-B DNA annotation was obtained from (Kouzine *et al*., 2017).

### Quantification and statistical analyses

Statistical analyses were performed using R version 3.3.1 to 3.6.1 (http://www.r-project.org). Repeatmasker was downloaded from UCSC genome browser on January 12, 2021.

## Acknowledgements

We thank members of the Keeney and Jasin laboratories for discussions. We thank the MSK Integrated Genomics Operation for sequencing. MSK core facilities are supported by NCI Cancer Center support grant P30 CA08748. This work was supported by NIH grant R35 GM118092 (to S.K.).

