## Supplemental material for "Triple-helix potential of the mouse genome"

**Supplemental table S1. Genome assembly versions used in this study**

| Species | Genome version |
|---|---|
| Mouse | mm10 |
| Rat | rn6 |
| Chinese hamster | criGriChoV2 |
| Kangaroo rat | dipOrd1 |
| Naked mole-rat | hetGla2 |
| Squirrel | speTri2 |
| Pika | ochPri3 |
| Malayan flying lemur | galVar1 |
| Tree shrew | tupBel1 |
| Human | hg38 |
| Yeast | sacCer2 |
| Mouse lemur | micMur2 |
| Tarsier | tarSyr2 |

### Supplemental Figure Legends

**Figure S1. Additional examples of SPO11-independent S1-seq clusters.**
(**A**) Strand-specific S1-seq (RPM) at another representative DSB hotspot (left side, coincident with a peak in the SPO11-oligo sequencing) with two reproducible SPO11-independent read clusters nearby (right side).

**Figure S2. Hoogsteen base pairs and S1-seq counts at pyrimidine repeats.**
(**A**) Hoogsteen base pairs ($C^{+}$GC and TAT) and reverse Hoogsteen base pairs (CGG, TAA, and TAT). H-y isomers are composed of Hoogsteen base pairs and H-r isomers are composed of reverse Hoogsteen base pairs. The dotted lines indicate hydrogen bonds; the gray rectangles highlight the bonds in Hoogsteen pairs.
(**B**) Comparison *Spo11*$^{-/-}$ S1-seq counts at individual polypyrimidine mirror and non-mirror repeats. Boxes indicate median and interquartile range; whiskers indicate the most extreme data points which are 1.5 times the interquartile range from the box; outliers are not shown.
(**C**) Mean *Spo11*$^{-/-}$ S1-seq read densities at various polypyrimidine mirror repeats.

**Figure S3. Incomplete digestion of ssDNA in H-DNA can account for observed S1-seq patterns at $C(TC)_{20}$ sequences.**
As in **Figure 3**, in all panels, gray arrowheads indicate which ssDNA segments are digested with nuclease S1, and the adaptors are color coded to indicate whether the resulting sequencing read will map to the pyrimidine strand (blue) or purine strand (red). At the bottom of each schematic, the expected mapping position and strand for the S1-seq read(s) are shown.
(**A**) Predicted S1-seq patterns if the central loop and orphan strand (but not junction strand) of each H-DNA isomer is digested with nuclease S1. In this scenario, all four isomers are predicted to yield a central read (pyrimidine strand for H-y5 and H-r3; purine strand for H-y3 and H-r5). In addition, digestion of H-r3 or H-y3 would leave a 5′ overhang that, after fill-in with T4 DNA polymerase and ligation to a sequencing adaptor, is predicted to yield a second central read mapping to the opposite strand. In contrast, digestion of H-y5 or H-r5 would yield a 3′ overhang, which if inefficiently polished would not yield any S1-seq read. If this scenario reflects the major S1 digestion pattern, it would explain why central reads are more abundant than junction reads.
(**B**) Predicted S1-seq patterns if nuclease S1 cleaves the junction strand, digests the orphan strand to leave variable end points, and fails entirely to digest the central loop. In this scenario, H-y3 and H-r3 would yield two DNA ends with 3′ overhangs. If these were inefficiently polished, no S1-seq read would result, as shown; if polished, they would yield reads indistinguishable from complete cleavage as shown in **Figure 3C**. In contrast, H-y5 and H-r5 would yield 5′ overhangs that, after fill-in with T4 DNA polymerase, would yield a junction read plus an opposite-strand read inside the mirror repeat. For H-y5, this would yield a pyrimidine-strand junction read and purine-strand reads within the left half of the mirror repeat, consistent with the striated S1-seq signal observed (**Figures 1G and 1H**).
(**C**) Chemical sensitivity of a plasmid-borne H-y5-forming sequence, $(TC)_{17}$ (Glover *et al.*, 1990).

**Figure S4. Additional examples of S1-seq patterns around mirror repeats**
(**A**) Unusual S1-seq read distribution at a pyrimidine mirror repeat with a 10-bp interruption between the repeats. In this example, roughly equal numbers of central reads were observed on both strands.
(**B**) S1-seq signal at $CCC(TCTCCC)_7$ (n=143). Note that the fine-scale spatial pattern is distinct from $C(TC)_{20}$ (**Figure 1H**), but is highly stereotyped across individual copies of this repeat.

**Figure S5. Purine-strand S1-seq maps around imperfect C(TC)$_n$ repeats of various lengths.**
(**A**) Averaged purine-strand S1-seq signal (read counts) for C(TC)$_n$ repeats of the indicated lengths. The left graphs show signal for perfect repeats (same repeats as shown in **Figure 4D**); the right graphs show signal for imperfect repeats (same repeats as shown in **Figure 4H**).
(**B**) Heat maps of purine-strand S1-seq maps for imperfect C(TC)$_{20}$, C(TC)$_{26}$, and C(TC)$_{30}$ repeats. These are the same repeats shown in **Figures 4E, 4F, and 4G**.

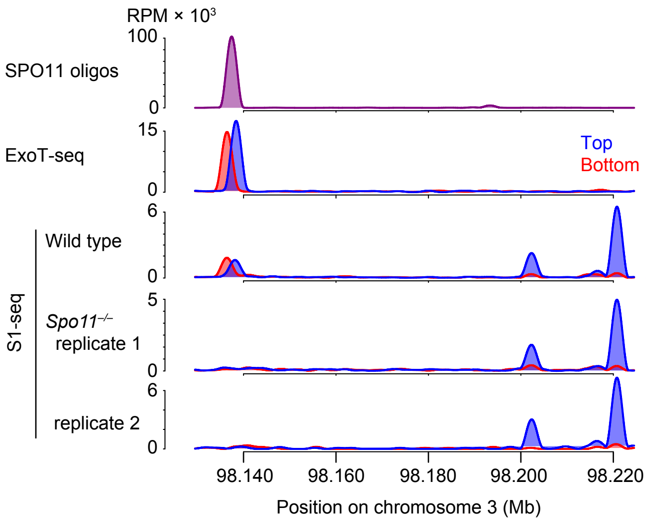

RPM × 10³
100
0
SPO11 oligos
15
0
ExoT-seq
Top
Bottom
6
0
Wild type
S1-seq
5
0
Spo11−/−
replicate 1
6
0
replicate 2
98.140
98.160
98.180
98.200
98.220
Position on chromosome 3 (Mb)

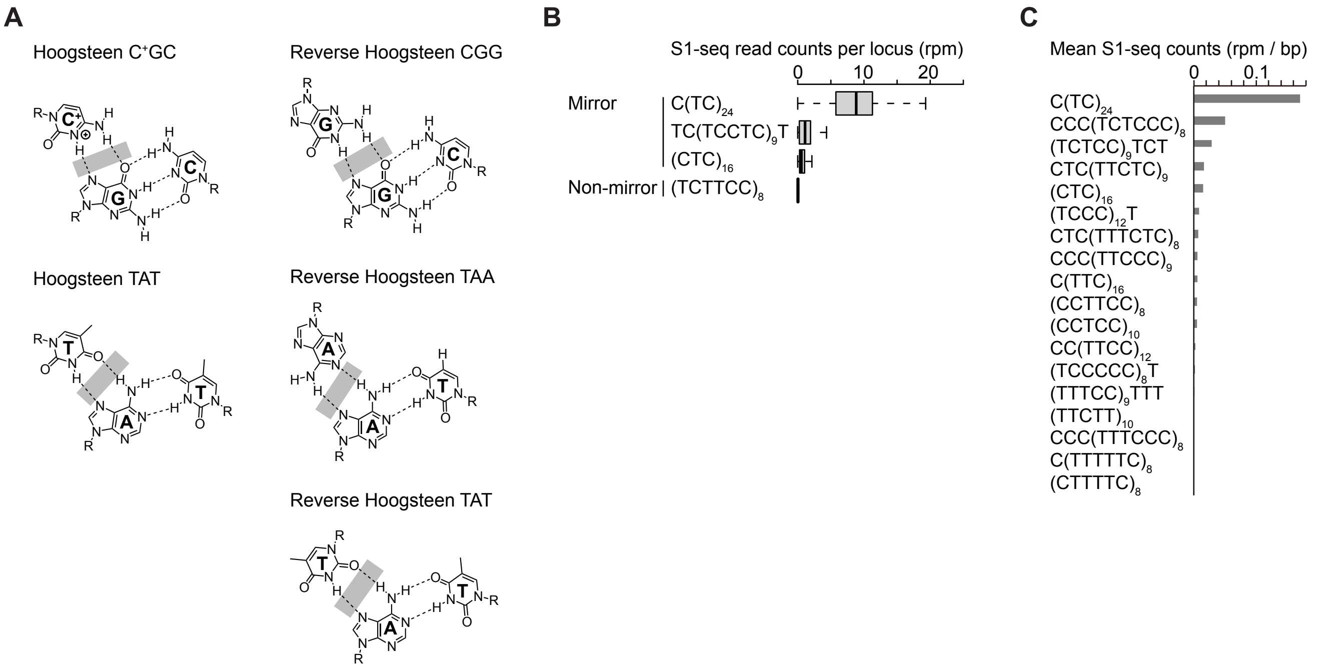
A
Hoogsteen C+GC
Reverse Hoogsteen CGG
Hoogsteen TAT
Reverse Hoogsteen TAA
Reverse Hoogsteen TAT
B
S1-seq read counts per locus (rpm)
0
10
20
Mirror
C(TC)24
TC(TCCTC)9T
(CTC)16
Non-mirror
(TCTTCC)8
C
Mean S1-seq counts (rpm / bp)
0
0.1
C(TC)24
CCC(TCTCCC)8
(TCTCC)9TCT
CTC(TTCTC)9
(CTC)16
(TCCC)12T
CTC(TTTCTC)8
CCC(TTCCC)9
C(TTC)16
(CCTTCC)8
(CCTCC)10
CC(TTCC)12
(TCCCCC)8T
(TTTCC)9TTT
(TTCTT)10
CCC(TTTCCC)8
C(TTTTTC)8
(CTTTTC)8

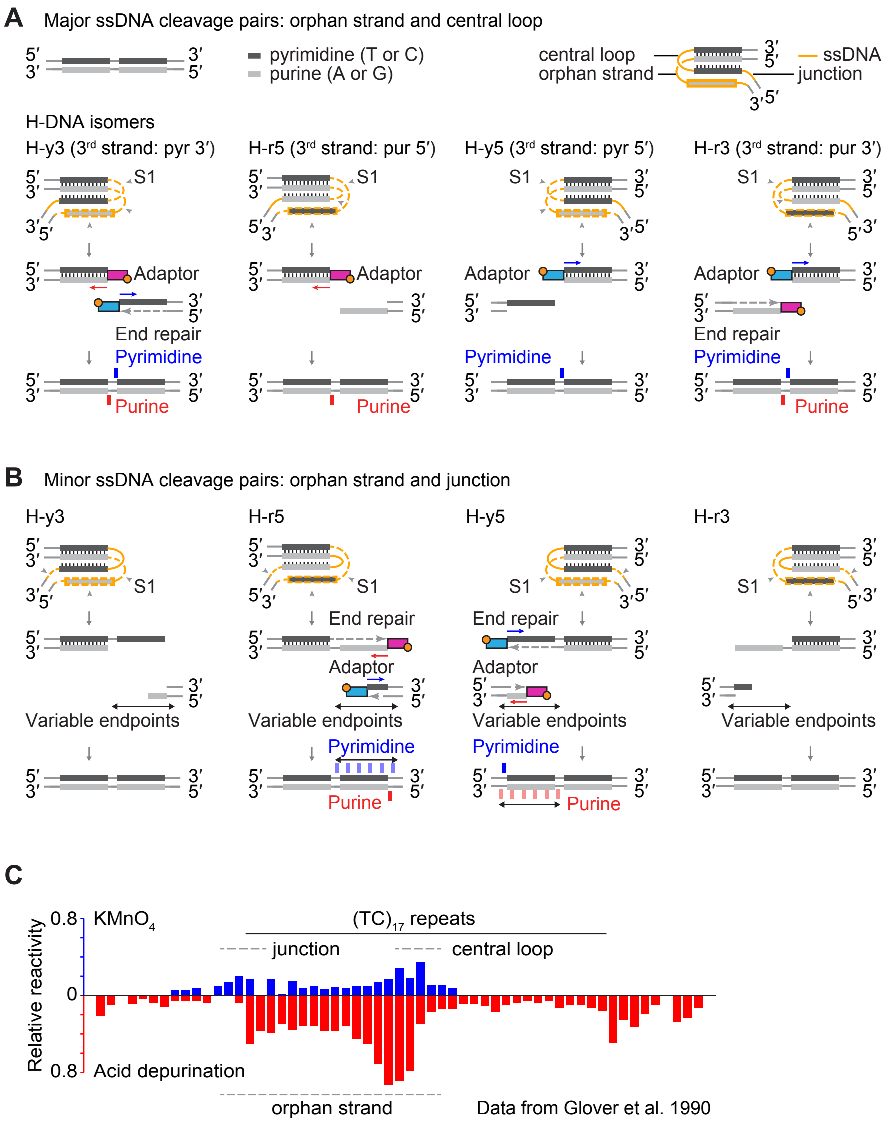
A Major ssDNA cleavage pairs: orphan strand and central loop
pyrimidine (T or C)
purine (A or G)
central loop
orphan strand
ssDNA
junction
H-DNA isomers
H-y3 (3rd strand: pyr 3′)
H-r5 (3rd strand: pur 5′)
H-y5 (3rd strand: pyr 5′)
H-r3 (3rd strand: pur 3′)
S1
Adaptor
End repair
Pyrimidine
Purine
B Minor ssDNA cleavage pairs: orphan strand and junction
H-y3
H-r5
H-y5
H-r3
Variable endpoints
C
Relative reactivity
KMnO4
Acid depurination
(TC)17 repeats
junction
central loop
orphan strand
Data from Glover et al. 1990

**A**

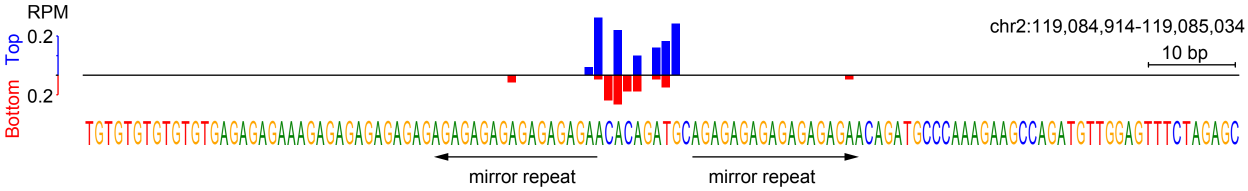
RPM
Top
Bottom
0.2
0.2
chr2:119,084,914-119,085,034
10 bp
TGTGTGTGTGTGTGAGAGAGAAAGAGAGAGAGAGAGAGAGAGAGAGAGAGAGAACACAGATGCAGAGAGAGAGAGAGAGAACAGATGCCCAAAGAAGCCAGATGTTGGAGTTTCTAGAGC
mirror repeat
mirror repeat

**B**

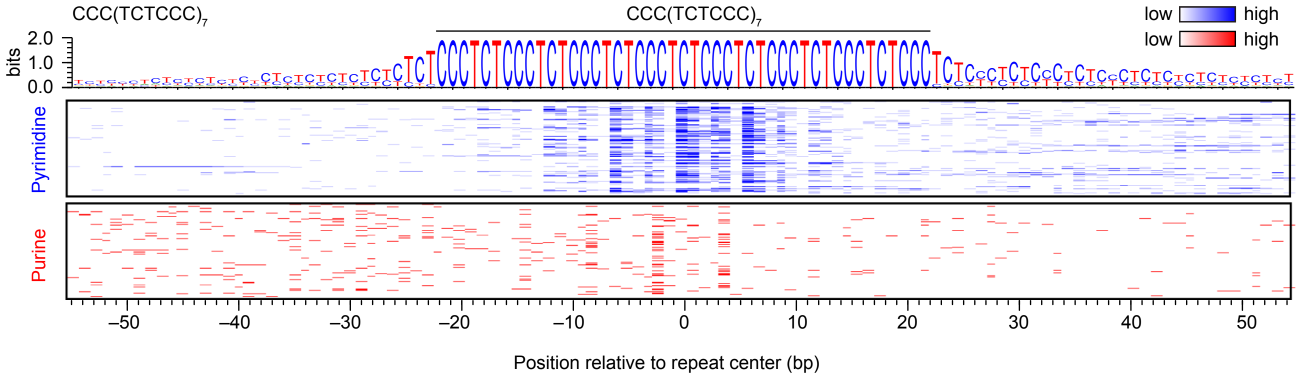
CCC(TCTCCC)7
CCC(TCTCCC)7
low
high
low
high
bits
2.0
1.0
0.0
Pyrimidine
Purine
−50
−40
−30
−20
−10
0
10
20
30
40
50
Position relative to repeat center (bp)

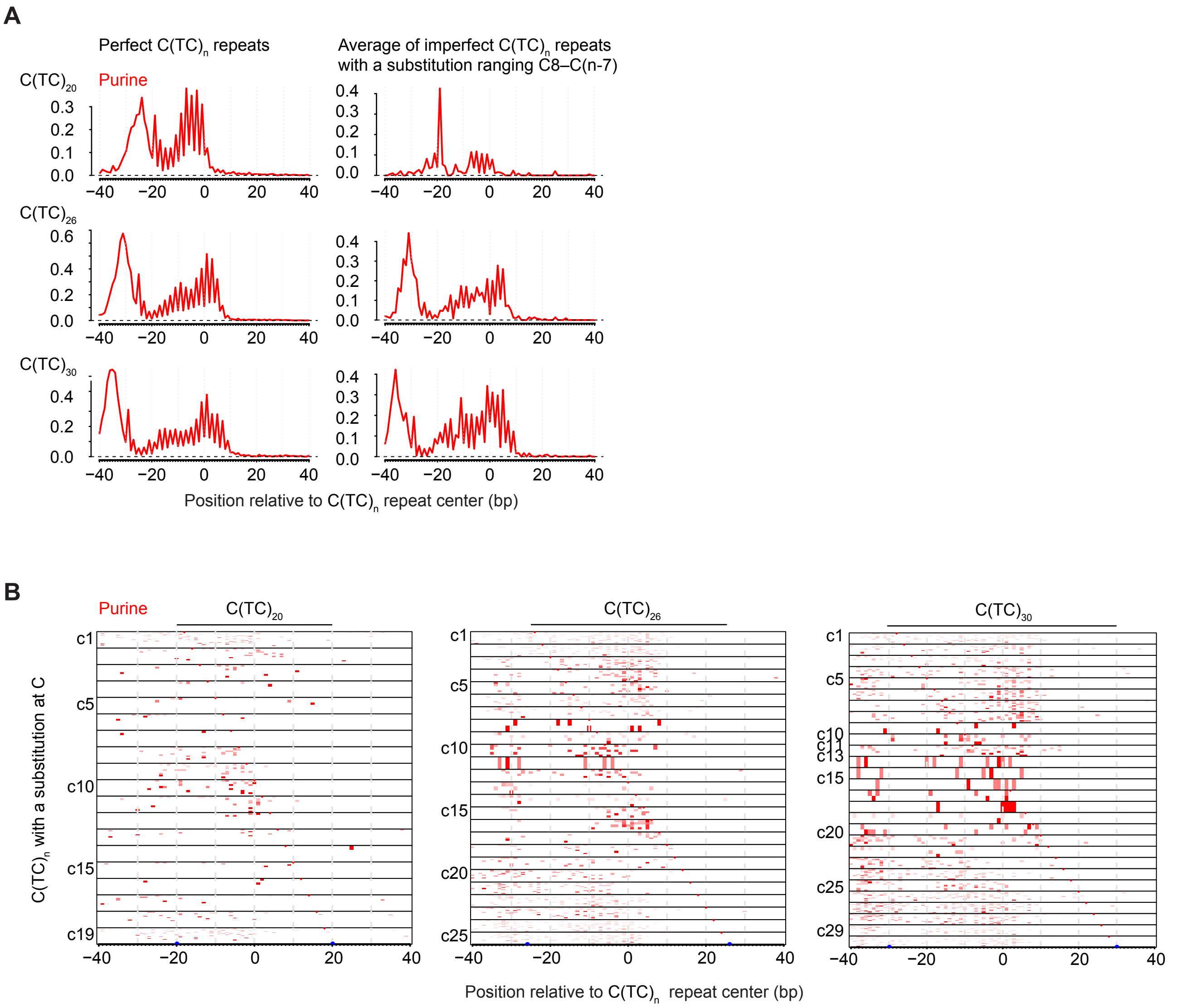
A
Perfect C(TC)n repeats
Average of imperfect C(TC)n repeats with a substitution ranging C8–C(n-7)
C(TC)20
Purine
C(TC)26
C(TC)30
Position relative to C(TC)n repeat center (bp)
B
Purine
C(TC)20
C(TC)26
C(TC)30
C(TC)n with a substitution at C
Position relative to C(TC)n repeat center (bp)
